# An orthogonal Cro/O_R_3 binary system for targeted gene expression

**DOI:** 10.64898/2025.12.24.696356

**Authors:** Kazuhide Asakawa, Kimiko Saka, Yukiko Yoshida, Saki Yamaura, Haruna Nakajo

## Abstract

Precise spatiotemporal control of gene expression is essential for dissecting complex biological systems. Here, we introduce a phage-derived Cro/O_R_3 binary system for transcriptional activation that functions robustly in zebrafish and human cells. A Cro repressor fused to the VP16 activation domain efficiently drives expression of O_R_3 operator-linked genes, and its activity can be targeted to specific cell types using enhancer-trap and promoter-controlled zebrafish lines. As a proof of concept, we manipulate glutamate transporter Glast/Slc1a3b-expressing cells in the central nervous system by Cro/O_R_3-mediated expression of a human ataxia-associated, uptake-deficient Glast/Slc1a3b variant, causing spastic, uncoordinated movement and axial deformation. Concurrent Gal4/UAS-mediated visualization reveals substantially disrupted positioning of motor neurons, highlighting their selective sensitivity to extracellular glutamate homeostasis during spinal circuit assembly. Together, these results establish the Cro/O_R_3 system as an orthogonal platform for targeted gene expression, enabling precise mechanistic dissection of how specific cell types and their interactions underlie diverse biological processes.

## Introduction

Multicellular organisms arise through the differentiation of cells and the coordination of their activities, enabling the emergence of complex physiological functions. The establishment of such elaborate organization critically depends on the precise execution of genetic programs and downstream cellular behaviors at the right time and place. Binary gene expression systems can be used not only to induce gene expression in cultured cells but also to drive expression in specific cell populations within multicellular organisms, allowing precise spatiotemporal control of gene activity. These systems typically rely on naturally occurring or engineered transcription factors that activate gene expression by binding to specific upstream elements often arranged in tandem repeats. When such a transcription factor is expressed in a genetically defined cell population, a gene of interest linked to the upstream elements is expressed accordingly, allowing targeted genetic manipulation of specific cell populations and evaluation of their physiological outcomes in vivo.

A single binary gene expression system can be applied across multiple species if the functional interaction between the transcription factor and its target DNA sequence occurs among diverse host cells, exemplified by the Gal4/UAS system ^1^, Q system ^2^, LexA/LexAop system ^3^, and Tet ON/OFF system ^4^. By leveraging the modular domain structure of transcription factors, the functionality of binary gene expression systems can be tailored through domain swapping to optimize them for the host organism. For example, the Gal4/UAS system originating from budding yeast ^5^ can stimulate transcription in heterologous host cells, including human cells ^6,7^, and has been extensively applied in multicellular model organisms to target gene expression to specific tissues or developmental stages ^1,8–10^. The DNA-binding domain of Gal4 can be replaced by another domain with different DNA-binding specificity to induce gene expression under the control of different DNA elements ^6^. Moreover, the transcription activation domain can be replaced with that of another transcription factor to modulate transcriptional activity ^11^, or with a chemically inducible domain that allows conditional activation of transcription through binding to a specific compound ^12^. The applicability of binary gene expression systems is further elevated by a comprehensive and well-characterized repertoire of readily usable driver, reporter, and effector lines. In this regard, *Drosophila* benefits from an extensive collection of genetic resources centered on the Gal4/UAS system ^13^. However, compared with invertebrate models, vertebrate systems remain limited by the incomplete availability of well-characterized driver lines, restricted compatibility among distinct binary systems, and a paucity of tools enabling parallel or intersectional gene manipulation at cellular resolution ^9,14–17^.

Here, we report the development of a binary expression system based on the bacteriophage λ repressor Cro, which controls the switch between lytic and lysogenic cycles (Fig. 1a). Bacteriophage λ encodes two repressor proteins, CI and Cro, both of which bind the same three operator sites (O_R_1, O_R_2, and O_R_3) within the right operator (O_R_) region of the λ genome ^18,19^. CI has a high affinity for O_R_1 and O_R_2 and promotes transcription for the maintenance of the lysogenic state. On the other hand, the repressor Cro exhibits a higher affinity for O_R_3 and promotes the lytic cycle by repressing CI and turning down early lytic promoters. We found that Cro can be converted into a transcriptional activator when attached to acidic transcriptional activation domains from VP16 (CroF2 and CroF3) and efficiently drove expression of genes adjacent to the O_R_3 operator repeats in zebrafish and human cells. We established a collection of promoter-controlled and enhancer trap CroF2 and CroF3 zebrafish driver lines. As a proof of concept, we ablated glutamate transporter Slc1a3b/Glast-expressing cells in the central nervous system (CNS) using the Cro/O_R_3 system, leading to an axial bending phenotype. Moreover, expression of an uptake-deficient glutamate transporter variant carrying a human ataxia– associated mutation recapitulated the axial phenotype, highlighting the importance of glutamate homeostasis in the CNS for axial straightening.

**Fig. 1.**
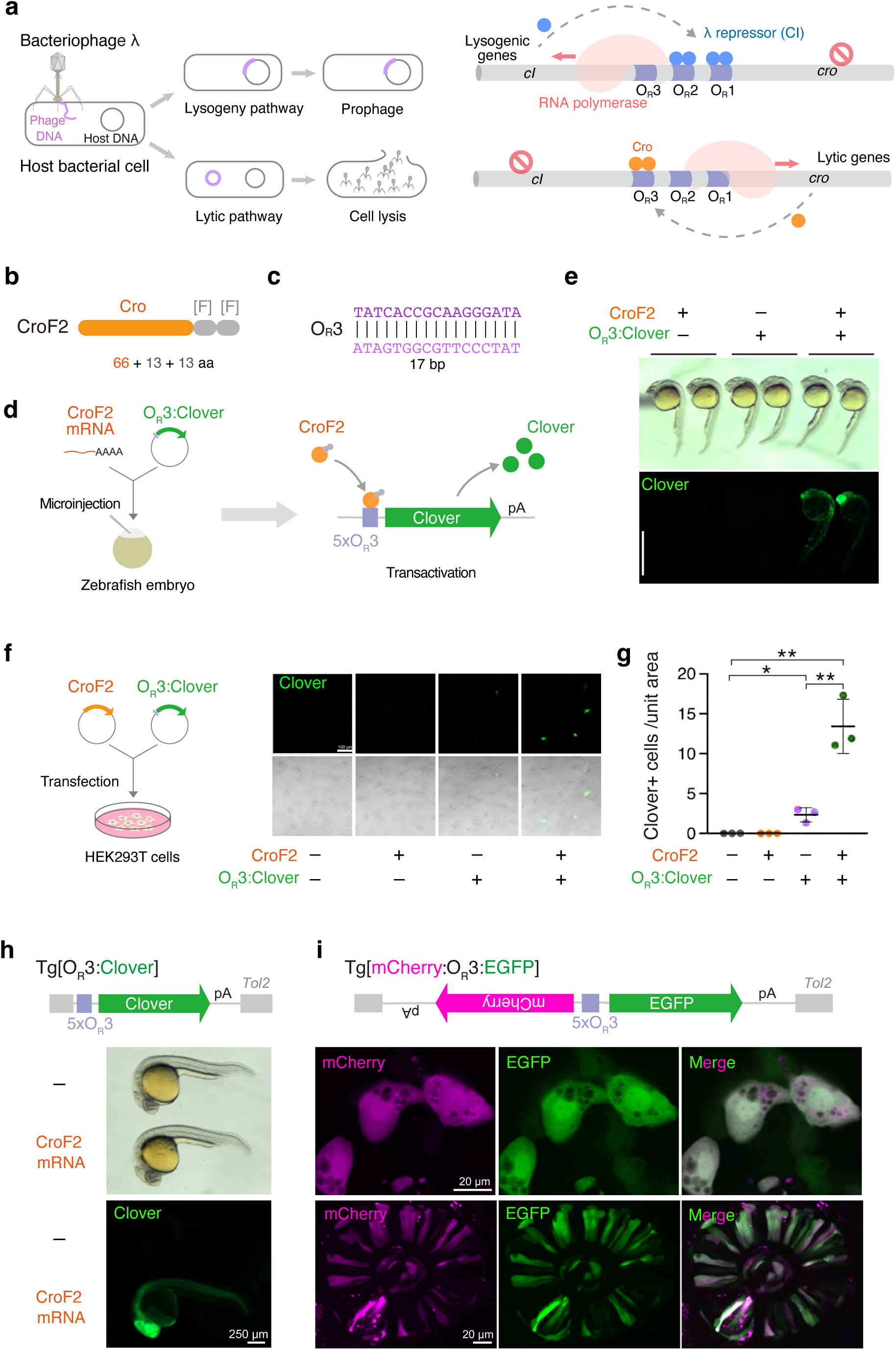
Cro/O_R_3-mediated gene expression in zebrafish and human cells (a) Lytic and lysogenic cycles of bacteriophage λ (left). CI maintains lysogeny by binding to O_R_1/O_R_2 to repress lytic promoters, whereas Cro binding to O_R_3 reduces CI expression to promote lytic development (right). (b) CroF2 consists of Cro (66 amino acids) fused to two F domains (26 amino acids). (c) The nucleotide sequence of O_R_3. (d) Co-injection of CroF2 mRNA and an O_R_3:Clover plasmid into one-cell-stage zebrafish embryos leads to Clover expression. (e) Cro-dependent Clover expression in embryos at 24 hpf. Scale bar, 1mm. (f) Cro/O_R_3-mediated transactivation in mammalian cells. HEK293T cells were co-transfected with CroF2 and O_R_3:Clover plasmids, resulting in CroF2-dependent Clover expression. (g) Quantification of Clover+ cells. The number of Clover+ cells was counted in each condition in three independent transfection experiments. *P=0.0102 (grey/purple), **P=0.0024 (grey/green), **P=0.0053 (purple/green) (t test, two-tailed). The center bars indicate the mean and the error bars show the standard deviation (SD). (h) Clover expression driven by Tg[O_R_3:Clover] upon CroF2 expressed by CroF2 mRNA injection at the one-cell stage. (i) Bidirectional expression of Clover and mCherry from Tg[mCherry:O_R_3:Clover] upon CroF2 expressed by CroF2 mRNA injection at the one-cell stage.

Independent Gal4/UAS-mediated visualization of spinal neurons uncovered a pronounced, non-cell-autonomous effect of glutamate dyshomeostasis on motor neuron positioning. Together, these results establish the Cro/O_R_3 system as an efficient and orthogonal platform for gene expression, enabling precise dissection of gene function in defined cell types.

## Results

### Cro/OR3 system: a binary gene expression system in zebrafish and human cells

Inspired by the DNA sequence-specific interaction between Cro and the O_R_3 operator in bacteriophage 𝜆 (Fig. 1a), we aimed to design a modular binary gene expression system based on the Cro/O_R_3 interaction that operates in vertebrate systems. To harness the Cro/O_R_3 interaction for transcriptional activation, we fused Cro with the two transcription activation modules from VP16 consisting of 13 amino acids containing a critical phenylalanine ([F] domain) ^20,21^ and designated it as CroF2 (Fig. 1b). To investigate whether CroF2 functions as a transcriptional activator in vertebrate cells, we injected in vitro-transcribed CroF2 mRNA into zebrafish embryos at the one-cell stage together with a reporter plasmid carrying the gene encoding the green fluorescent protein Clover ^22^ under the control of the five tandem O_R_3 repeats (O_R_3:Clover) (Fig. 1c, d). In the injected embryos at 24 hours post-fertilization (hpf), mosaic Clover signals were observed, but not in control embryos injected with the O_R_3:Clover plasmid alone (Fig.1e), indicating CroF2-dependent Clover expression in zebrafish. To further assess the applicability of the Cro/O_R_3 system in mammalian cells, we co-transfected plasmids encoding CroF2 and O_R_3:Clover into HEK293T cells. Robust Clover signals were detected in the cells co-transfected with CroF2 and O_R_3:Clover, whereas non-transfected control and cells transfected with CroF2 alone showed no Clover expression (Fig. 1f, g). Cells transfected with O_R_3:Clover alone showed some CroF2-independent Clover expression, although the number of Clover-positive cells was markedly lower than the co-transfected condition. Taken together, these observations demonstrate that CroF2 drives Clover expression via the O_R_3 repeats and that the Cro/O_R_3 system functions as a binary gene expression system in zebrafish and human cells.

To test whether CroF2 can drive gene expression from a genomically integrated O_R_3 transgene, we generated a transgenic zebrafish carrying a stable O_R_3:Clover transgene (Tg[O_R_3:Clover]). Injection of CroF2 mRNA into Tg[O_R_3:Clover] embryos at the one-cell stage gave rise to ubiquitous Clover expression at 24 hpf (Fig. 1h). We then investigated whether the Cro/O_R_3 system could direct bidirectional gene expression as does the Gal4/UAS system in the regulation of *GAL1*/*GAL10* promoter of the yeast *Saccharomyces cerevisiae* ^5^. An O_R_3 transgene in which mCherry and EGFP genes were placed in opposite directions on both sides of the five tandem O_R_3 repeats was generated (Tg[mCherry:O_R_3:EGFP]) (Fig. 1i). Injection of CroF2 mRNA into the zebrafish embryos carrying Tg[mCherry:O_R_3:EGFP] at the one-cell stage resulted in the expression of both mCherry and EGFP in various tissues, including the epithelial cells and retinal neural progenitor cells at 24 hpf (Fig. 1i). These observations show that the Cro/O_R_3 system operates as a bidirectional gene expression system. We noted that, in the absence of CroF2, a transient, weak Clover signal was occasionally detectable in the caudal epithelium of Tg[O_R_3:Clover] fish during early embryonic stages (Sup. Fig. 1), suggesting that the 5xO_R_3 repeats can exhibit a low level of basal promoter activity depending on cell type and developmental stage.

### Targeted gene expression by the Cro/O_R_3 system

The key advantage of binary gene expression systems lies in their ability to achieve precise spatiotemporal control of gene expression in multicellular organisms. To examine whether the Cro/O_R_3 system enables reliable targeted gene expression in vivo, we expressed CroF2 in zebrafish spinal motor neurons (SMNs) using a bacterial artificial chromosome (BAC) transgenic line in which the CroF2 gene was inserted into the *mnr2b* locus (Tg[mnr2b-hs:CroF2]) (Fig. 2a) ^23–25^. In parallel, we generated a reporter line carrying an O_R_3-driven mScarlet-tagged nitroreductase transgene (Tg[O_R_3:mScarlet-NTR]) (Fig. 2b). In Tg[mnr2b-hs:CroF2] Tg[O_R_3:mScarlet-NTR] double-transgenic larvae, robust mScarlet-NTR expression was observed in SMNs (Fig. 2c). To validate the identity of these CroF2-driven mScarlet-NTR–expressing cells, we compared their labeling with a pan–motor neuron EGFP reporter expressed through the Gal4/UAS system under control of the same mnr2b-BAC^25^. Nearly all mScarlet-NTR–positive cells coexpressed EGFP at 48 hpf (Fig. 2d), indicating that CroF2-driven O_R_3 activation faithfully recapitulates SMN-specific expression. Together, these results demonstrate that the Cro/O_R_3 system enables reliable cell type-specific gene expression in zebrafish.

**Fig. 2.**
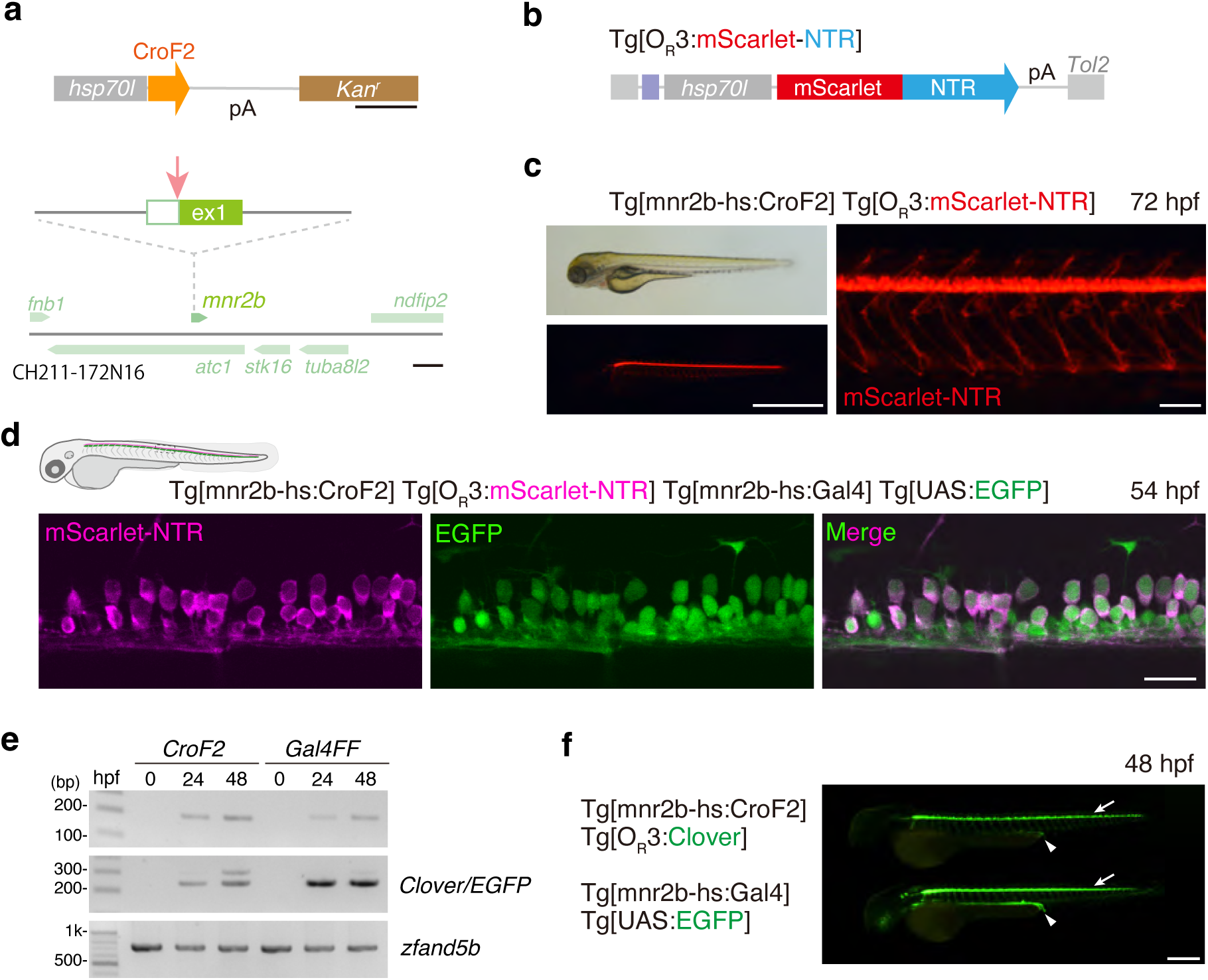
Targeted gene expression by the Cro/O_R_3 system (a) Construction of Tg[mnr2b-hs:CroF2]. The *hsp70l:CroF2* cassette was introduced just in front of the start codon of *mnr2b* gene in the BAC (CH211-172N16). Scale bars, 500 bp (top), 10 kb (bottom). (b) The structure of Tg[O_R_3:mScarlet-NTR] (c) A lateral view of Tg[mnr2b-hs:CroF2] Tg[O_R_3:mScarlet-NTR] fish at 3 dpf. Scale bars, 1 mm (left) and 100 µm (right). (d) A lateral view of the spinal cord of Tg[mnr2b-hs:CroF2] Tg[O_R_3:mScarlet-NTR] Tg[mnr2b-hs:Gal4] Tg[UAS:EGFP] fish at 54 hpf. The dashed lines demarcate the ventral limit of the spinal cord. Scale bar, 20 µm. (e) Expression levels of CroF2/Gal4FF (top), Clover/EGFP (middle), and zfand5b (bottom) transcripts in Tg[mnr2b-hs:CroF2];Tg[O_R_3:Clover] and Tg[mnr2b-hs:Gal4];Tg[UAS:EGFP] fish. The results shown are representative of three independent experiments. bp, base pairs. (f) A lateral view of Tg[mnr2b-hs:CroF2];Tg[O_R_3:Clover] (top) and Tg[mnr2b-hs:Gal4];Tg[UAS:EGFP] (bottom) fish at 48 hpf. The arrows and arrowheads indicate the spinal motor columns and intestines, respectively, highlighted by green fluorescence. Scale bar, 500 µm.

The parallel activation of O_R_3- and UAS-linked reporters by CroF2 and Gal4FF expressed from the same mnr2b-BAC prompted us to directly compare the transcriptional output of the two systems. We analyzed Tg[mnr2b-hs:CroF2]; Tg[OR3:Clover] and Tg[mnr2b-hs:Gal4]; Tg[UAS:EGFP] larvae, in which both transcription factors have two [F] domains for transcription activation, are driven by identical *mnr2b* regulatory elements and control reporters linked to five tandem binding sites. RT-PCR analysis revealed comparable transcript levels of CroF2 and Gal4FF at both 24 and 48 hpf (Fig. 2e). In contrast, the transcript level of Clover was lower than that of EGFP, consistent with weaker green fluorescence in CroF2-expressing SMNs (Fig. 2f). These findings indicate that both the CroF2/O_R_3 and Gal4FF/UAS systems support robust, targeted gene expression in vivo, although the CroF2/O_R_3 system exhibits slightly lower transcriptional activation than Gal4FF/UAS under otherwise equivalent conditions.

### Targeting CroF2 expression by enhancer trapping

Enhancer trapping provides a powerful approach for labeling cell populations that cannot be readily targeted using known *cis*-regulatory elements. To extend the utility of the Cro/O_R_3 system, we performed an enhancer trap screen using a *Tol2* transposon-based cassette in which the CroF2 gene was placed under the control of the *hsp70l* promoter (*hsp70l*:CroF2) (Fig. 3a). Fish injected with the enhancer trap cassette were crossed with homozygous Tg[O_R_3:Clover] fish (Fig. 3b). Germline insertions of the enhancer trap cassette into genomic loci subject to enhancer regulation resulted in Clover expression in the F1 fish. In total, we identified 100 independent Clover expression patterns among F1 fish obtained from 301 injected fish (33%). As an example, to assess tissue-specific CroF2 expression, we examined the CroF2-driven Clover-labeled cells in the head region by using the Tg[elavl3:mCherry] line as a reference for brain tissue^26^. The CroF2 trap lines drove Clover expression in discrete clusters of cells in specific brain regions, including the forebrain, diencephalon (habenula), midbrain (tectum), and hindbrain, demonstrating the effectiveness of enhancer trapping for isolating CroF2 driver lines that label defined cell populations (Fig. 3c). We reasoned that incorporating an additional [F] domain into CroF2 at its carboxy-terminus (yielding CroF3) might be tolerated by zebrafish cells and amplify Clover expression from Tg[O_R_3:Clover] reporter. Based on this rationale, we engineered CroF3 and performed an enhancer-trap screening (Fig. 3d). Our ongoing screening using the *hsp70l:CroF3* cassette yielded multiple CroF3 driver lines (10 independent Clover expression patterns in F1 out of 27 injected fish: 33%), including a specific driver line for peripheral sensory organs (Fig. 3e, f). Clover expression patterns driven by CroF2 or CroF3 in F1 fish identified through ongoing enhancer trap screening are continuously updated and made available in the NIG-Zebra database (https://nig-zebra.nbrp.jp/image/).

**Fig. 3.**
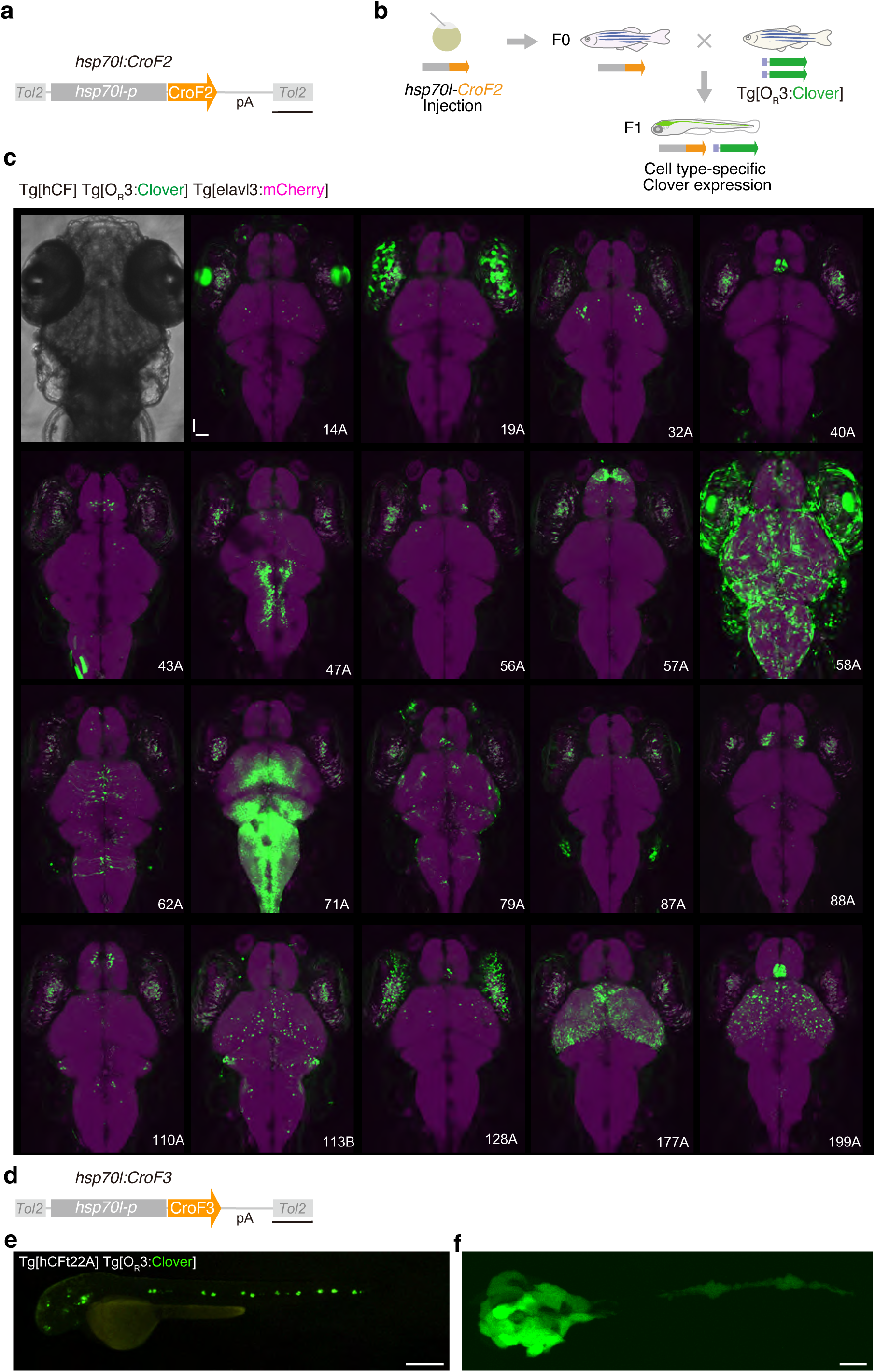
Enhancer trap screening of tissue- and cell type–specific CroF2 driver lines (a) Structure of the *Tol2* transposon-based CroF2 enhancer-trap construct. Scale bar, 200 bp. (b) Scheme of enhancer trap screening. Fish injected with the enhancer trap cassette and *Tol2* transposase mRNA at the one-cell stage were raised to adulthood and crossed with homozygous Tg[O_R_3:Clover] fish. (b) Representative examples of distinct enhancer-driven CroF2 expression patterns in the head region at 5-6 dpf, visualized by Clover expression from the Tg[O_R_3:Clover] reporter. The whole brain structure was highlighted in magenta by Tg[elavl3:mCherry]. Major Clover expression sites of the enhancer-trap lines were as follows: 14A (lens), 19A (retinal pigment epithelium), 32A (optic tectum), 40A (pineal gland), 43A (forebrain), 47A (hindbrain), 56A (habenula), 57A (forebrain), 58A (lens, epithelium), 62A (radial cells in midbrain and hindbrain), 71A (optic tectum, hindbrain), 79A (olfactory epithelium, epithelial layer overlying the brain), 87A (otic vesicle), 88A (habenula), 110A (forebrain), 113B (optic tectum, cerebellum), 128A (retina), 177A (optic tectum), 199A (optic tectum, pineal gland). All larvae exhibited a green autofluorescence signal from the retinal epithelium. Scale bar, 50 µm. (d) Structure of the *Tol2* transposon-based CroF3 enhancer-trap construct. Scale bar, 200 bp. (e) A lateral view of Tg[hCFt22A] Tg[O_R_3:Clover] at 48 hpf. Tg[hCFt22A] driver express CroF3 in peripheral sensory organs, including neuromasts. Scale bar, 500 µm (f) A confocal z-stack image of a neuromast in Tg[hCFt22A] Tg[O_R_3:Clover] fish at 48 hpf. Scale bar, 10 µm

### Ablation of Glast/Slc1a3b-expressing cells results in axial bending

To extend the utility of the Cro/O_R_3 system from cell visualization to functional analysis, we evaluated its capacity to manipulate physiological processes via targeted gene expression. As a test case, we focused on glutamate transporters, which mediate both glutamate uptake and efflux, regulate diverse cellular functions, and maintain excitatory amino acid homeostasis across tissues. In the CNS, the glutamate transporters expressed in glial cells play a critical role in regulating excitatory neurotransmission and preventing glutamate-induced excitotoxicity by rapidly clearing glutamate from the synaptic cleft ^27^.

Given this critical role of glutamate transporters in excitatory neurotransmission, we used the Cro/O_R_3 system to examine the contribution of Slc1a3b/Glast, a glutamate transporter predominantly expressed in astrocytes ^28–30^, to glutamate homeostasis in the CNS. A stable transgenic zebrafish line, Tg[slc1a3b:CroF2], in which CroF2 is driven by the ∼9.5-k base-pair sequence upstream of the zebrafish *slc1a3b* translational start site ^30^ was established (Fig. 4a). When combined with Tg[O_R_3:mScarlet-NTR], Tg[slc1a3b:CroF2] drove mScarlet-NTR expression broadly in the CNS (Fig. 4b, c). Consistent with the predominant expression of *slc1a3b* in astrocytes ^30^, sparse labeling through injecting the O_R_3:Clover plasmid at the two-cell stage labeled CroF2-expressing astrocytes in the CNS with Clover at 3 dpf (Fig. 4c). We also noted that some SMNs and interneurons were also labeled, suggesting that *slc1a3b* is also expressed in a population of spinal neurons (Supplementary Fig. 2). Treatment of Tg[slc1a3b:CroF2] Tg[O_R_3:mScarlet-NTR] fish with 10 mM Mtz from 24 hpf for 48 hours resulted in a substantial reduction of the mScarlet signal in the CNS (Fig. 4f), confirming the ablation of mScarlet-NTR-expressing cells.

**Fig. 4.**
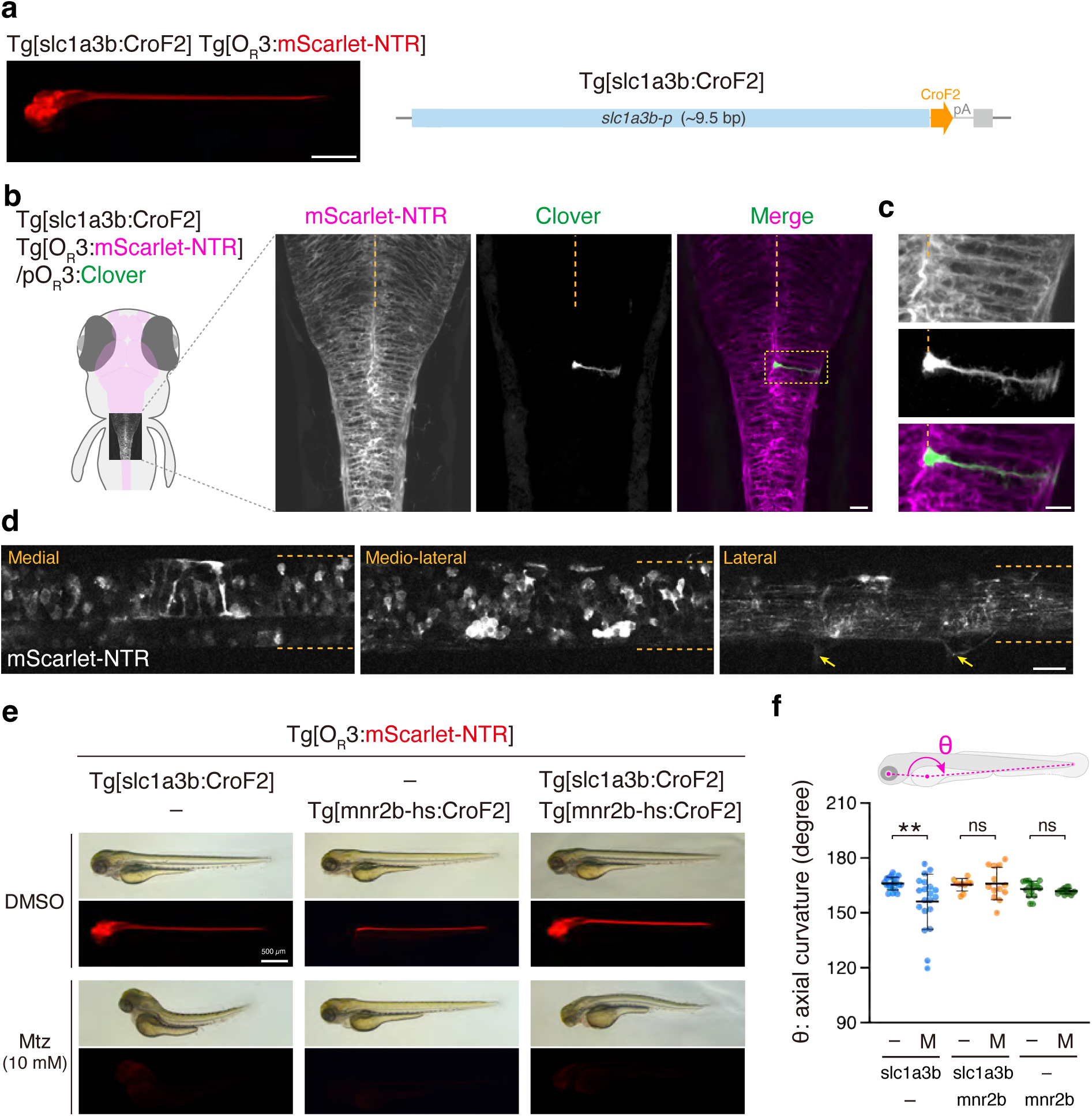
NTR-mediated ablation of *slc1a3b*-expressing cells by the Cro/O_R_3 system (a) Tg[slc1a3b:CroF2] Tg[O_R_3:mScarlet-NTR] fish at 3 dpf (left). Scale bar, 500 µm. The structure of Tg[slc1a3b:CroF2] (right). The CroF2 gene is linked to the ∼9.5-kb sequence upstream of the zebrafish *slc1a3b* translational start site. (b) Single confocal sections of the spinal cord of Tg[slc1a3b:CroF2] Tg[O_R_3:mScarlet-NTR] fish at 3 dpf at distinct levels of the mediolateral axis (lateral views, the dashed outline in panel b). Arrows indicate motor axons. The dashed lines demarcate the dorsal and ventral limits of the spinal cord. Scale bar, 20 µm. (c) A dorsal view of hindbrain-spinal cord junction in Tg[slc1a3b:CroF2] Tg[O_R_3:mScarlet-NTR] fish at 3 dpf, which was injected with the OR3:Clover plasmid at the 2-cell stage. Dashed lines indicate midline. Scale bar, 20 µm. (d) A single astrocyte labeled with Clover in panel b. Scale bar, 10 µm. (e) Chemogenetic ablation of NTR-expressing cells in Tg[scl1a3b:CroF2] Tg[O_R_3:mScarlet-NTR], Tg[mnr2b-hs:CroF2] Tg[O_R_3:mScarlet-NTR], and Tg[slc1a3b:CroF2] Tg[mnr2b-hs:CroF2] Tg[O_R_3:mScarlet-NTR] fish at 3 dpf. Fish were treated with DMSO or 10 mM metronidazole (Mtz) at 24 hpf for 48 hours. (f) Axial curvature (θ), the angle formed between the line connecting the lens center to the yolk center and the line connecting the yolk center to the tail tip, was measured. Twenty Tg[slc1a3b:CroF2] Tg[O_R_3:mScarlet-NTR] fish from five independent crosses were examined for each of – (DMSO) and Mtz conditions. Tg[slc1a3b:CroF2] Tg[mnr2b-hs:CroF2] Tg[O_R_3:mScarlet-NTR] fish from three independent crosses were examined for each of – (DMSO, ten fish) and Mtz (thirteen fish) conditions. Fifteen Tg[mnr2b-hs:CroF2] Tg[O_R_3:mScarlet-NTR] fish from three independent crosses were examined for each of – (DMSO) and Mtz conditions. **P = 0.0073 (Mann-Whitney test, two-tailed). ns, not statistically significant. The error bars show SD.

Intriguingly, besides the reduction of mScarlet-NTR signals, Tg[slc1a3b:CroF2] Tg[O_R_3:mScarlet-NTR] larvae also exhibited an abnormal body curvature (Fig. 4f, g). The axial bending phenotype was not observed after ablation of *mnr2b*-positive SMNs by treatment with 10 mM Mtz in Tg[mnr2b-hs:CroF2]; Tg[O_R_3:mScarlet-NTR] larvae (Fig. 4f, g). Furthermore, co-ablation of *slc1a3b*-expressing SMNs and *mnr2b*-expressing cells significantly, if not fully, suppressed the axial bending phenotype in Tg[slc1a3b:CroF2] Tg[mnr2b:CroF2] Tg[O_R_3:mScarlet-NTR] larvae treated with 10 mM Mtz (Fig. 4f, g). Together, these observations suggest that ablation of *slc1a3b*-expressing cells induces axial bending at least through an SMN-dependent mechanism and highlight a critical role for glutamate homeostasis in the CNS in maintaining axial straightening.

### Glast/Slc1a3b-mediated glutamate homeostasis is essential for spinal motor circuit assembly

One potential cause of the axial bending phenotype induced by ablation of *slc1a3b*-expressing cells is reduced Slc1a3b-dependent glutamate uptake, leading to extracellular glutamate dyshomeostasis in the CNS. To test whether impaired glutamate uptake alone is sufficient to elicit this phenotype and to probe the underlying intercellular communication, we expressed a mutant Slc1a3b protein carrying the P283R substitution, which corresponds to the P290R mutation in human GLAST/SLC1A3 identified in patients with episodic ataxia type 6 ^31^. This mutation markedly reduces glutamate uptake capacity and exerts a dominant-negative effect through multimerization with the wild-type transporter ^31^. For cell-type-specific expression, we generated a transgenic O_R_3 line (Tg[O_R_3:slc1a3b-P283R]) that co-expresses Slc1a3b-P283R and mScarlet via a self-cleaving T2A peptide (Fig. 5a) and crossed it with the Tg[slc1a3b:CroF2] driver line. Among the offspring, larvae expressing mScarlet in the CNS exhibited an axial bending phenotype at 3 dpf (Fig. 5b, c), recapitulating the phenotype observed upon *slc1a3b*-cell ablation. At 5 dpf and beyond, Tg[slc1a3b:CroF2] Tg[O_R_3:slc1a3b-P283R] larvae continued to show abnormal axial curvature, and frequently displayed intermittent episodes of spastic, uncoordinated swimming movements, likely driven by unbalanced, excessive muscle contractions (Supplementary Movie 1, 2), and eventually died. Together, these observations indicate that impaired Slc1a3b-dependent glutamate uptake compromises motor function.

**Fig. 5.**
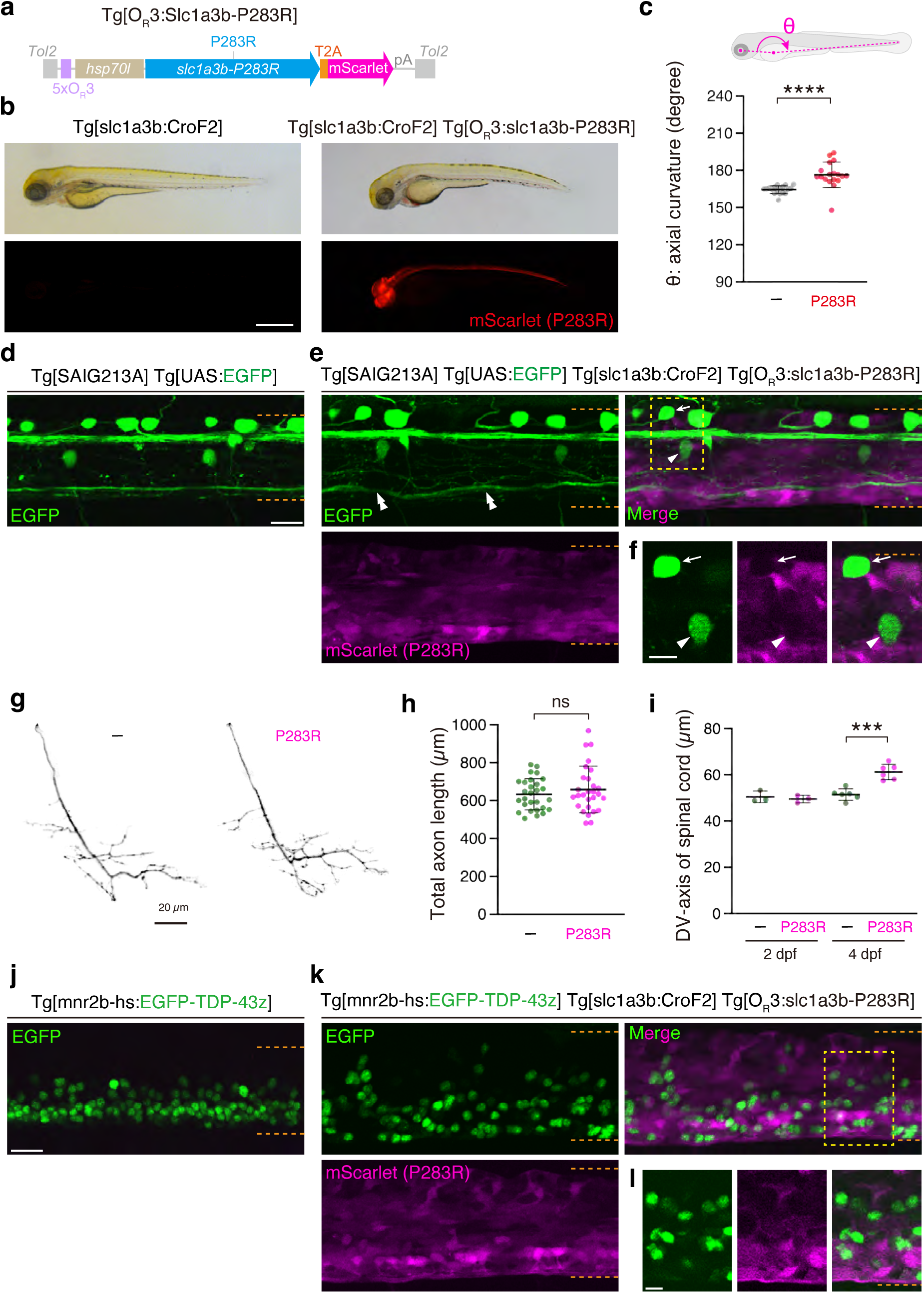
Glutamate homeostasis influences SMN positioning (a) The structure of Tg[O_R_3:slc1a3b-P283R], which expresses Slc1a3b-P283R and mScarlet via the self-cleaving T2A peptide. (b) Tg[slc1a3b:CroF2] and Tg[slc1a3b:CroF2] Tg[O_R_3:slc1a3b-P283R] fish at 3 dpf (right). Scale bar, 500 µm. (c) Axial curvature (θ), the angle formed between the line connecting the lens center to the yolk center and the line connecting the yolk center to the tail tip, was measured in each condition in panel b. Eighteen fish obtained from 4 independent crossed were analyzed in each condition. ****P < 0.0001 (Mann-Whitney test, two-tailed). The error bars indicate SD. (d) A lateral view of the spinal cord in Tg[SAIG213A] Tg[UAS:EGFP] fish at 4 dpf. The dashed lines demarcate the dorsal and ventral limits of the spinal cord. Scale bar, 20 µm. (e) A lateral view of the spinal cord in Tg[SAIG213A] Tg[UAS:EGFP] Tg[slc1a3b:CroF2] Tg[O_R_3:slc1a3b-P283R] fish at 4 dpf. The arrow, arrowhead and double-arrowhead indicate a Rohon-Beard cell, a CaP and axonal tracts running along the ventral spinal cord, respectively. (f) A single confocal section of the region indicated by the dashed square in panel d. Scale bar, 10 µm. (g) Representative motor axons of CaP in Tg[SAIG213A] Tg[UAS:EGFP] (–) and Tg[SAIG213A] Tg[UAS:EGFP] Tg[slc1a3b:CroF2] Tg[O_R_3:slc1a3b-P283R] (P283R) fish at 48 hpf. Scale bar, 20 µm. (h) Total lengths of the CaP axons. Data were obtained from seven (mScarlet-negative siblings, 28 cells) and seven (mScarlet-positive fish, 28 cells) animals. ns, not statistically significant (P = 0.3706, t-test, two-tailed). The error bars indicate SD. (i) Lengths of the dorso-ventral axis of the spinal cords. Data were obtained from three (mScarlet– siblings) and three (mScarlet+ fish) animals at 2 dpf, and six (mScarlet– siblings) and six (mScarlet+ fish) animals at 4 dpf. ***P = 0.0002, t-test, two-tailed). The error bars indicate SD. (j) A lateral view of the spinal cord in Tg[mnr2b-hs:EGFP-TDP-43z] fish at 4 dpf. The dashed lines demarcate the dorsal and ventral limit of the spinal cord. Scale bar, 20 µm. (k) A lateral view of the spinal cord in Tg[mnr2b-hs:EGFP-TDP-43z] Tg[slc1a3b:CroF2] Tg[O_R_3:slc1a3b-P283R] fish at 4 dpf. (l) A single confocal section of the region indicated by the dashed square in panel k. Scale bar, 10 µm.

Given the dependence of Slc1a3b-mediated axial straightening on SMNs (Fig. 4g), we next asked how glutamate homeostasis influences the assembly of spinal neural circuits. To visualize spinal neurons under the Slc1a3b-P283R-induced conditions, we employed the Gal4 driver Tg[SAIG213A] and the reporter Tg[UAS:EGFP], which in combination label spinal neurons, including Rohon-Beard sensory neurons and caudal primary motor neurons (CaPs) ^32^. This orthogonal visualization showed that EGFP-labeled RB cells and CaPs were mScarlet-negative (i.e., *slc1a3b*-negative) and that their positions appeared largely normal within the spinal cord of Tg[slc1a3b:CroF2] Tg[O_R_3:slc1a3b-P283R] Tg[SAIG213A] Tg[UAS:EGFP] quadruple transgenic fish at 4 dpf, in comparison to the Tg[SAIG213A] Tg[UAS:EGFP] double transgenic fish (Fig. 5d, e, f). Consistently, the trajectories and total lengths of CaP axons were not significantly different between these two conditions (Fig. 5g, h). In contrast to this preserved cellular pattern, the spinal cord itself was noticeably thicker at 4 dpf, likely reflecting mechanical compression associated with excessive muscle contraction and axial bending (Fig. 5e, i). Moreover, the EGFP-labeled axonal tracts running along the ventral spinal cord exhibited distorted trajectories (Fig. 5e), raising the possibility that the ventral spinal cord is particularly susceptible to Slc1a3b-P283R-induced glutamate dyshomeostasis.

To investigate the organization of the ventral spinal cord more closely, we visualized the spatial distribution of SMN somata by labeling their nuclei with EGFP-tagged zebrafish TDP-43/Tardbp (hereafter, EGFP-TDP-43z), a predominantly nuclear RNA/DNA-binding protein, expressed from the BAC transgene Tg[mnr2b-hs:EGFP-TDP-43z] ^33^. In zebrafish, SMNs are tightly clustered along the entire length of the ventral spinal cord, and accordingly, their EGFP-TDP-43z-labeled nuclei appeared densely aligned in this region (Fig. 5j). However, under Slc1a3b-P283R–induced conditions at 4 dpf, the EGFP-labeled motor neuron nuclei, which were mScarlet-negative (i.e., Slc1a3b-P283R– negative), were more dispersed, with some nuclei abnormally positioned in the dorsal spinal cord in Tg[slc1a3b:CroF2] Tg[O_R_3:slc1a3b-P283R] Tg[mnr2b-hs:EGFP-TDP-43z] larvae (Fig. 5k). This phenotype indicates that Slc1a3b-P283R–induced glutamate dyshomeostasis disrupts SMN distribution in the spinal cord. Because most SMNs were mScarlet-negative (Fig. 5l), the disrupted SMN distribution was likely caused in a non-cell-autonomous manner.

Altogether, these findings demonstrate that Slc1a3b-mediated glutamate homeostasis is essential for proper spatial organization of SMNs and, consequently, for the assembly of functional spinal motor circuits. Disruption of glutamate uptake in Slc1a3b-expressing cells perturbs SMN positioning in a non–cell-autonomous manner, revealing a critical role for extracellular glutamate homeostasis in shaping spinal circuit architecture. Moreover, these results highlight the utility of the Cro/O_R_3 system as an orthogonal genetic tool to interrogate multi–cell-type interactions in vivo through targeted gene expression in combination with existing genetic approaches.

## Discussion

Understanding gene and cellular functions in multicellular organisms requires technologies that allow precise spatiotemporal control of gene expression. In the present study, we report the development of a phage-derived Cro/O_R_3 binary gene expression system applicable in zebrafish and human cells. We demonstrated that the expression of the Cro repressor fused to the VP16 activation domain led to robust expression of O_R_3-linked transgenes.

Furthermore, this transactivation can be confined to specific cell populations using promoter/enhancer-driven CroF2 and CroF3 driver fish lines. We further extended the range of cell types accessible through the Cro/O_R_3 system by performing enhancer trapping and successfully established 101 lines in which diverse cell types were labeled with CroF2 and CroF3. Because the Cro/O_R_3 system operates independently of existing binary systems such as Gal4/UAS, Q, and LexA, it provides orthogonal control. When used together with these systems, it enables independent manipulation of multiple cell populations within a single animal, meeting the emerging need to investigate inter–cell type and inter-organ communication. Further expansion of the Cro/O_R_3 system will require the generation of additional tissue- and cell-type specific driver lines using enhancer trapping and genome editing-mediated knock-in, as well as of versatile O_R_3 lines for cellular- and subcellular imaging and manipulation. In parallel, the CI/O_R_1 and CI/O_R_2 repressor systems for the lysogenic cycle of λ phage warrant exploration as candidates for extending the repertoire of targeted gene expression systems, with careful evaluation of their potential cross-interference with the Cro/O_R_3 system. From a broader perspective, it will also be important to determine whether this system functions reliably across diverse species other than zebrafish and humans, thereby validating its general applicability.

Efficient binary gene expression requires optimization in several key aspects. We found that the present Cro/O_R_3 system based on CroF2 and 5xO_R_3 repeats is slightly lower than the Gal4FF/UAS system in terms of transcriptional activation, despite their shared use of two [F] domains as the activation domain. Although it remains unclear whether this difference reflects an intrinsic property of Cro itself, arises from the interaction between Cro and the 5xO_R_3 repeats, or involves other factors, these observations indicate that the present Cro/O_R_3 system still has room for improvement. The modular architecture of CroF2 allows modulation of transcriptional activity by appending or removing the [F] domain ^21^, when targeting cell populations with weak or strong promoters, respectively. Indeed, as part of our ongoing work, we are continuing enhancer-trap screens with Cro fused to three [F] domains (CroF3), which has enabled more frequent isolation of driver lines, presumably through enhanced transcriptional amplification that facilitates detection of weak enhancers. Another consideration is that, in zebrafish, UAS transgenes often undergo progressive transcriptional silencing over generations ^34^, and this silencing is correlated with DNA methylation of the multicopy UAS. While we have confirmed over four generations that *mnr2b* BAC-driven CroF2 yields stable and consistent Clover expression from Tg[O_R_3:Clover] in SMNs, whether O_R_3-transgene silencing might occur in future generations, and to what extent, will need to be carefully monitored. Taking this point into account, determining the optimal copy number of O_R_3 repeats for gene expression, as well as exploring potential optimization of O_R_3 sequence, will be important directions for future refinement and improvement.

One of our primary motivations for developing this technique was to establish Cro/O_R_3 as a system capable of inducing dominant phenotypes in vivo, thereby enabling functional assessment of genes and cell types. First, we demonstrated that Cro/O_R_3-mediated chemogenetic ablation of Slc1a3b-expressing cells resulted in an axial bending phenotype. This phenotype was attenuated by concurrent ablation of *mnr2b*-expressing motor neurons, indicating that axial straightening is achieved, at least in part, through an SMN-dependent mechanism. A similar axial bending phenotype has been reported in fish treated with epinephrine or injected with Urp1/Urotensin neuropeptide ^35^, underscoring the contribution of an adrenergic cerebrospinal fluid-contacting neuron–Urotensin axis to axial straightening. How impaired Slc1a3b-dependent glutamate homeostasis affects these adrenergic neurons, and thereby modulates the SMN-dependent output, remains an open question and represents an opportunity to dissect the cellular logic of axial straightening.

Second, building on the cell-ablation results, Cro/O_R_3-mediated expression of an uptake-deficient Slc1a3b-P283R variant recapitulated the axial deformation, supporting the notion that impaired Slc1a3b-dependent glutamate clearance, and the resulting extracellular glutamate dyshomeostasis, drives this phenotype. While single-cell analyses provided little evidence that this dyshomeostasis perturbs motor axon outgrowth, its impact on the spatial organization of SMNs was substantial. Because most SMNs did not express Slc1a3b-P283R, their disrupted positioning likely reflects a non–cell-autonomous effect arising from elevated extracellular glutamate due to halted astrocytic uptake ^30^. This controlled perturbation offers a tractable means to interrogate astrocyte-derived excitotoxicity at single-cell resolution in the intact CNS and to define how glutamate imbalance shapes spinal circuit assembly and motor output. Insights into how Slc1a3b-P283R–induced dyshomeostasis affects SMNs may also help uncover the cellular basis of episodic ataxia 6 ^31^ and other neurological disorders involving glutamate excitotoxicity, including amyotrophic lateral sclerosis (ALS) ^36^. Moreover, this system provides a powerful platform to explore activity-dependent aspects of motor-neuron development ^37^.

In summary, this study shows that the Cro/O_R_3 system provides a versatile and robust platform for gene expression in zebrafish and human cells, enabling both functional interrogation of genes and cells. As demonstrated, applications that combine the Cro/O_R_3 system with other binary systems will enhance our capacity to investigate intercellular communication and organismal physiology.

## Methods

### Animals

This study was performed in accordance with the Guide for the Care and Use of Laboratory Animals of the Institutional Animal Care and Use Committee (IACUC) of National Institute of Genetics (NIG, approval numbers: R7-26). All experimental protocols were approved by the IACUC of NIG. All authors complied with the ARRIVE guidelines. Wild-type and transgenic zebrafish, hybrids of the AB and Tübingen strains, were used in all experiments. The fish were raised at 28°C under a 12-hour light/12-hour dark cycle (L/D) for the first five days post-fertilization. At these developmental stages, sex is not yet determined. Plasmids and transgenic zebrafish lines used in this study were described in detail in Supplementary information.

### RT-PCR

The total RNA was extracted from Tg[mnr2b-hs:CroF2] Tg[OR3:Clover] and Tg[mnr2b-hs:Gal4] Tg[UAS:EGFP] embryos at 0 hpf (25 or 50 embryos), 24 hpf (10-50 embryos) and 48 hpf (10-50 embryos) as previously described ^38^. One μg of the total RNA was used for cDNA synthesis using oligo dT (SuperScript™ IV First-Strand Synthesis System, Invitrogen, 18091050). PCR was performed using TaKaRa Ex Taq (Takara Bio Inc., RR001B). Cro and Gal4 transcripts were detected using a primer pair against the common 3’UTR sequence: 3’UTR_F (5′-CAT TAC CAA CTT GTC TGG TGT C-3′) and 3’UTR_R (5′-TGT AAC CAT TAT AAG CTG C-3′). Clover and EGFP transcripts were detected using a primer pair: Clover_GFP_F (5′-AAG GAG GAC GGC AAC ATC C T-3′) and Clover_GFP_R (5′-CTT CTC GTT GGG GTC TTT GC-3′). zfand5b transcripts were detected using a primer pair: zfand5b–133f (5′-ATA GTA CAC ACC GAA ACG GAC AC-3′) and zfand5b-772r (5′-TTA TAT TCT CTG GAT TTT ATC GGC-3′). Amplification was performed at an initial denaturation at 95 ℃ for 2 min, followed by 25 cycles of 15 s at 95 °C; 30 s at 55 °C (3’UTR_F and 3’UTR_R, zfand5b–133f and zfand5b–772r) or 60 °C (Clover_GFP_F and Clover_GFP_R); and 30 s at 72 °C, and a final extension at 72 °C for 5 min. PCR products were detected following electrophoresis on 2.0% agarose gels, and were imaged using Printgraph Classic (ATTO, WSE5400).

#### Culture cells and transfection

HEK293T cells were cultured in Dulbecco’s Modified Eagle Medium (DMEM; Nacalai tesque) supplemented with 10% fetal bovine serum (FBS) at 37 °C in a humidified incubator with 5% CO_2_. Cells were transfected with plasmid DNAs using Lipofectamine 3000 transfection reagent (Thermo Fisher Scientific) according to the manufacturer’s instructions.

#### Immunohistochemistry and microscopic analyses

HEK293T cells were seeded onto poly-L-lysine coated 4-well chamber slides (Matsunami glass) 24 hours post transfection. Cells were fixed with 4% paraformaldehyde, permeabilized with 0.2% Triton X-100 in PBS (Phosphate buffered saline), and blocked with 5% normal goat serum (NGS) in PBS. Cells were then incubated with anti-GFP antibody (1:2000; Aves labs, GFP-1010, AB_2307313) diluted in PBS containing 1% NGS for 1 hour at room temperature, washed with PBS, and incubated with anti-chicken Alexa Fluor 488 (1:500; Invitrogen) diluted in PBS containing 1% NGS for 1 hour at room temperature. After washing, samples were mounted with DAPI-Fluoromount-G (Southern Biotech). Images were acquired using Olympus FV3000 confocal microscope with 10x objective lens. Images were analyzed using ImageJ software (version 1.54p).

For confocal analyses of zebrafish, images were acquired from live fish embedded in 0.8–1% low-melting agarose (NuSieve GTG Agarose, Lonza) on a Glass Base dish (IWAKI, 3010-035) using an Olympus FV1200 laser confocal microscope with a 20x water immersion objective (NA1.0). The fish were raised in an embryonic buffer containing 0.003% (w/v) N-phenylthiourea (SIGMA, P7629) to inhibit melanogenesis. The hemicord at the level of spinal segments 13–16 was identified using the cloaca, typically located around the boundary between segments 16 and 17 along the rostrocaudal axis, as a marker. Confocal image stacks were acquired as serial optical sections along the z-axis, analyzed using Olympus Fluoview Viewer (Version 2.1b) and ImageJ (Version 2.16.0/1.54p), and processed for presentation using Adobe Photoshop (Version 22.5.9) and Illustrator (Version 27.0). The axon lengths were measured by Imaris Filament Tracer (Imaris 9.2.0). Stereoscope images were acquired using LEICA M165FC equipped with a K7 Color CMOS Microscope Camera.

#### Metronidazole (Mtz) treatment

Tg[slc1a3b:CroF2] Tg[O_R_3-mScarlet-NTR], Tg[mnr2b-hs:CroF2] Tg[O_R_3-mScarlet-NTR], or Tg[slc1a3b:CroF2] Tg[mnr2b-hs:CroF2] Tg[O_R_3-mScarlet-NTR] embryos were dechorionated and treated with freshly prepared 10 mM Mtz (SIGMA, M3761) in 0.2% DMSO in E3 buffer containing 0.003% (w/v) Phenylthiocarbamide (PTU) (SIGMA, P7629) from 24 to 72 hpf at 28 °C in the dark. The Mtz solution was replaced with a freshly prepared solution every 24 hours. For controls, the embryos were incubated in 0.2% DMSO in E3 buffer containing 0.003% PTU and mScarlet-negative embryos were treated with 10 mM Mtz in 0.2% DMSO in E3 buffer containing 0.003% PTU.

#### Enhancer Trapping

Wild type fish were injected with the pT2Zhsp70lCroF2 or pT2Zhsp70lCroF3 plasmid the synthesized mRNA encoding codon-optimized *Tol2* transposase ^39^ at the one-cell stage, raised and crossed with the homozygous Tg[O_R_3:Clover] reporter line.

## Data availability

Raw confocal imaging data (in .oib and .tif formats) of CroF2 enhancer trap lines have been deposited in figshare. Information on the expression patterns of additional CroF2 and CroF3 driver lines is available at the NIG-Zebra database (https://nig-zebra.nbrp.jp/image/), and the corresponding zebrafish lines are available from the authors upon request. All other data supporting the findings of this study are available within the article and Supplementary Information files. Source data are provided with this paper.

## Acknowledgement

The authors are grateful to Dr Fumihito Ono for providing Tg[elavl3:mCherry] line and to Kyoko Watanabe for technical assistance. This research was supported by KAKENHI Grant number JP 23K24219 (K.A.), "Strategic Research Projects " grant from ROIS (Research Organization of Information and Systems) (K.A.) and The National BioResource Project (NBRP) of the MEXT, Japan.

## Author contributions

K.A. conceived and designed the research. K.A., K.S., H.N., Y.Y., and S.Y. performed experiments and analyzed the data. K.A. wrote the manuscript. All authors discussed the results and commented on the manuscript.

## Supplementary Figures

**Supplementary Fig. 1.**
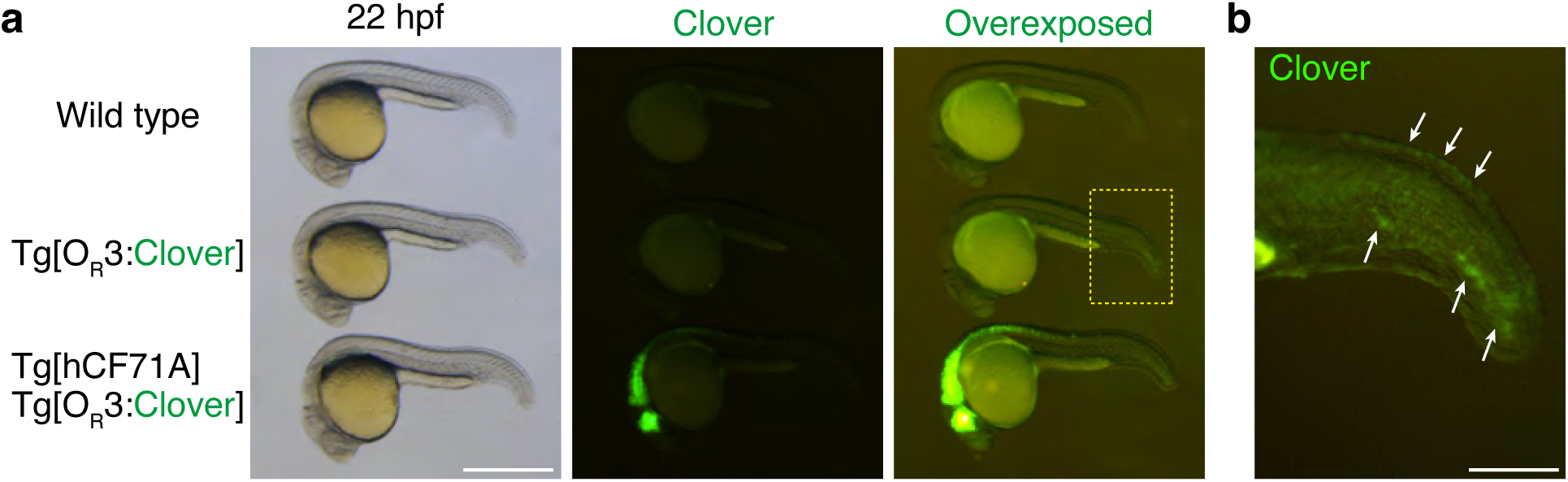
(a) A basal level expression of Clover from the Tg[O_R_3:Clover] transgene. A weak Clover signal detected in the caudal epithelium of Tg[O_R_3:Clover] fish during at 22 hpf (middle). Non-transgenic fish (Wild type) and Tg[hCF71A] Tg[O_R_3:Clover] fish, which is an enhancer trap line, were shown as negative and positive controls, respectively. Scale bar, 1 mm. (b) A magnified view of the dashed box in pane a. Arrow indicates Clover signals. Scale bar, 250 µm.

**Supplementary Fig. 2.**
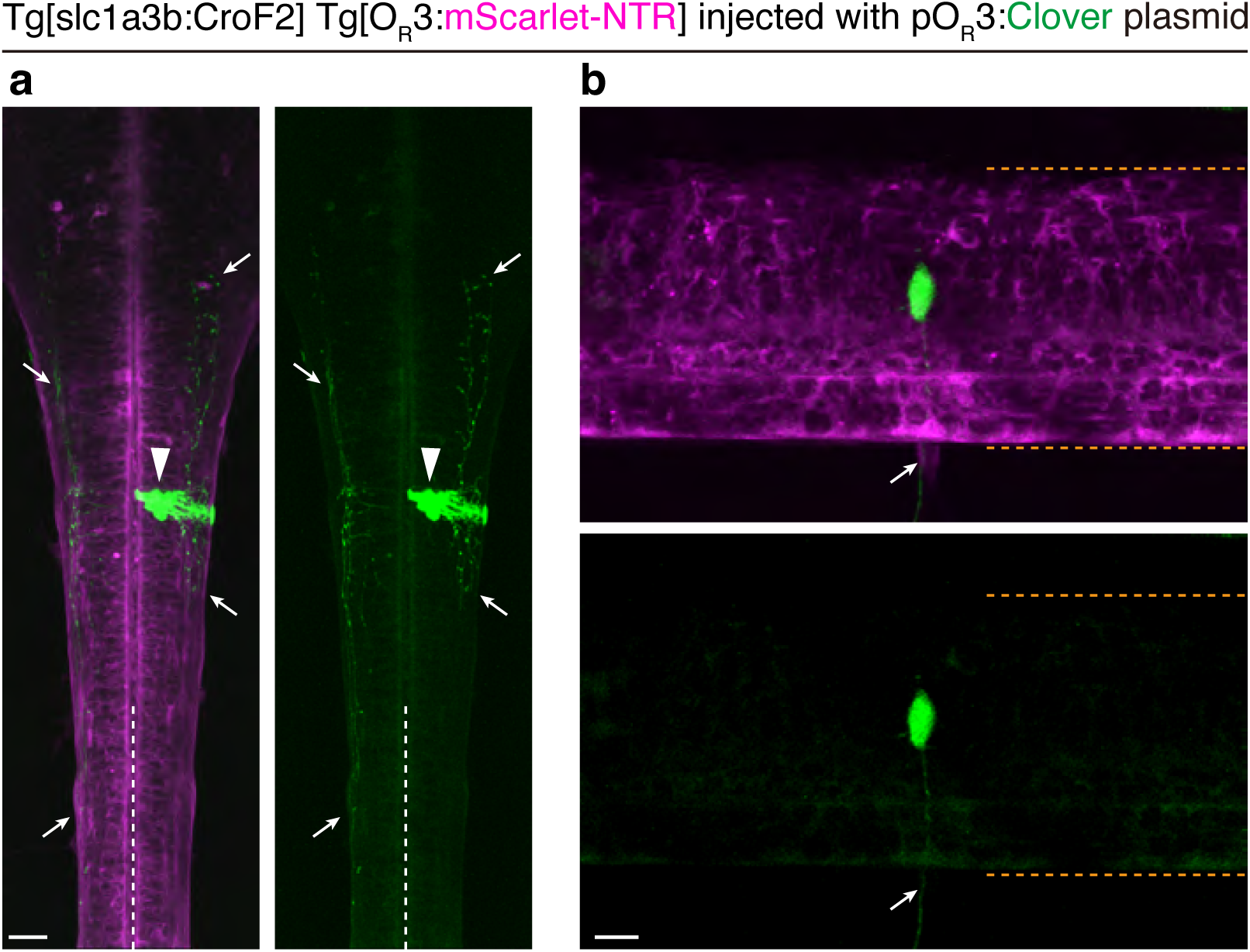
Sparse labeling of *slc1a3b*-expressing cells in the CNS. (a) Dorsal view of the hindbrain–spinal cord junction of Tg[slc1a3b:CroF2] Tg[O_R_3:mScarlet-NTR] larvae that were injected with the O_R_3:Clover plasmid at the two-cell stage. A cluster of Clover-expressing cells in the right hemicord (arrowhead), which are likely clonal, extends ipsilateral and contralateral axons (arrows). Dashed lines indicate the midline. Rostral is toward the top. Scale bar, 20 µm. (b) Dorsal view of the spinal cord. The arrow indicates a Clover-labeled motor axon exiting the spinal cord. Dashed lines demarcate the dorsal and ventral boundaries of the spinal cord. Rostral is to the right. Scale bar, 10 µm.

## Legends for Supplementary Movies

**Supplementary Movie1, and 2**

Swimming movements of Tg[slc1a3b:CroF2] Tg[O_R_3:slc1a3b-P283R] larvae. Offspring from the genetic cross between Tg[slc1a3b:CroF2] fish and Tg[O_R_3:slc1a3b-P283R] fish were sorted based on the red fluorescence of Slc1a3b-P283R. Slc1a3b-P283R-negative fish displayed normal free swimming behavior (Suppl. Movie1), while Slc1a3b-P283R-positive sibling showed abnormal swimming (Suppl. Movie2).

## Supplementary information

### Resource availability

Contact: Kazuhide Asakawa

Materials availability: Kazuhide Asakawa

Data availability: Kazuhide Asakawa

## Methods

The list of transgenic zebrafish lines used in this study

Tg[O_R_3:Clover] (this study)

Tg[mCherry:O_R_3:EGFP] (this study)

Tg[mnr2b-hs:CroF2] (this study)

Tg[O_R_3:mScarlet-NTR] (this study)

Tg[mnr2b-hs:Gal4] ^1^

Tg[UAS:EGFP] ^2^

Tg[elavl3:mCherry] ^3^

Tg[slc1a3b:CroF2] (this study)

Tg[O_R_3:slc1a3b-P283R] (this study)

Tg[mnr2b-hs:EGFP-TDP-43z] ^4^

The list of enhancer trap CroF2 lines established in this study is provided in Supplementary Information 2. Confocal z-section images of these CroF2 lines have been deposited in figshare. Expression pattern data for additional CroF2 lines will be made publicly available on the NIG-Zebra database (https://nig-zebra.nbrp.jp/image/) as they become available.

The plasmids used in this study

### pCS2CroF2

The DNA fragment carrying SP6 promoter-CroF2-SV40pA cassette was synthesized by Eurofins. CroF2 mRNA was produced by mMESSAGE mMACHINE™ SP6 Transcription Kit.

### pT2Zhsp70lCroF2 and pT2Zhsp70lCroF3

The *Tol2* transposon cassette carrying the *hsp70l* promoter (∼650 bp), CroF2 or *CroF3* gene, and SV40pA sequence was synthesized (GeneArt Gene Synthesis, Thermo Fisher Scientific).

### pcDNA3.1/CroF2 and pMA/5xO_R_3:Clover

These plasmids encoding codon-optimized CroF2 and Clover sequences for human expression, respectively, were generated using GeneArt gene synthesis (Thermo Fisher Scientific).

The plasmids used to generate the transgenic lines are described in the following transgenic lines section.

### Generation of transgenic zebrafish lines

All transgenic lines were generated by *Tol2* transposon-mediated transgenesis^5^. Details of the plasmids used for generating each transgenic line are summarized below.

Tg[O_R_3:Clover]

The *Tol2* transposon cassette carrying the 5xO_R_3, Clover gene, and SV40 polyadenylation sequence was synthesized (GeneArt Gene Synthesis, Thermo Fisher Scientific). The resulting pT2ZOR3Clover plasmid was used to generate Tg[O_R_3:Clover].

Tg[mCherry:O_R_3:EGFP]

The *Tol2* transposon cassette carrying five tandem O_R_3 sites, with mCherry linked to an *actb2* polyadenylation sequence and EGFP linked to an SV40 polyadenylation sequence arranged in opposite orientations, was synthesized (GeneArt Gene Synthesis, Thermo Fisher Scientific). The resulting pT2ZmCOR3G plasmid was used to generate Tg[mCherry:O_R_3:EGFP]

Tg[mnr2b-hs:CroF2]

Tg[mnr2b-hs:Gal4] was constructed as described in ^6^ except that the *hsp70l* promoter (650 bp)-CroF2-polyA-Km^r^ cassette was introduced downstream of the *mnr2b* 5’UTR in the mnr2b-BAC DNA (CH211-172N16). BAC engineering and *Tol2*-mediated transgenesis was performed as described in ^6^.

Tg[O_R_3:mScarlet-NTR]

The *Tol2* transposon cassette carrying the 5xO_R_3, mScarlet–fused NfsBzf1 gene ^7,8^, and SV40 polyadenylation sequence was synthesized (GeneArt Gene Synthesis, Thermo Fisher Scientific). The resulting pT2ZOR3mSNTR plasmid was used to generate Tg[O_R_3:mScarlet-NTR].

Tg[slc1a3b:CroF2]

The upstream promoter region of *slc1a3b* (9,550 bp) was amplified from the genomic DNA by PCR with the primer pair slc1a3b_p_f (5’-cacaaagcggcatccctgcggacc-3’) and slc1a3b_p_r (5’-gctgtcgctccacacactgagatc-3’). The cloned *slc1a3b* promoter was substituted for the *hsp70l* promoter in pT2Zhsp70lCroF2 to create the plasmid pT2Zslc1a3bCroF2, which was used to generate Tg[slc1a3b:CroF2].

Tg[O_R_3:Slc1a3b-P283R]

*Tol2* transposon cassettes carrying the 5xO_R_3 sequence, the *hsp70l* promoter the *slc1a3b-P283R* gene linked to mScarlet via a T2A peptide sequence, and the SV40 polyadenylation sequence were synthesized. (GeneArt Gene Synthesis, Thermo Fisher Scientific). The resulting pT2ZOR3hslc1a3bP283RT2AS was used to generate Tg[O_R_3:slc1a3b-P283R].

---

| Line name | Dissecting MS image (NIG-Zebra) | Major OR3:Clover expression sites | Head z-series, d5-d6 (.oib / .tif) |
| --- | --- | --- | --- |
| hCF14B | O | eye, otic vesicle, fin | O |
| hCF19A | O | eye | O |
| hCF32A | O | midbrain | O |
| hCF40A | O | guts, liver, gill, pinel gland | O |
| hCF43A | O | brain | O |
| hCF47A | O | hindbrain, spinal cord | O |
| hCF56A | O | brain | O |
| hCF57A | O | forebrain | O |
| hCF58A | O | lens, gill arches/jaw, notochord, som | O |
| hCF62A | O | brain, spinal cord | O |
| hCF71A | O | midbrain, hindbrain, spinal cord | O |
| hCF79A | O | mouth | O |
| hCF87A | O | otic vesicle, gill | O |
| hCF88A | O | brain, habenula | O |
| hCF110A | O | retina, forebrain | O |
| hCF113B | O | brain, somite | O |
| hCF128A | O | retina | O |
| hCF177A | O | midbrain, eye, somite | O |
| hCF199A | O | midbrain, spinal cord | O |

